# Utilizing Social Media and Video Games to Control #DIY Microscopes

**DOI:** 10.1101/053470

**Authors:** Maxime Leblanc-Latour, Craig Bryan, Andrew Pelling

**Author notes:** Author for correspondence: Andrew E. Pelling, 150 Louis Pasteur, MacDonald Hall, University of Ottawa, Ottawa, ON K1N 6N5, Canada, Web: http://www.pellinglab.net.

## Abstract

Open-source lab equipment is becoming more widespread with the popularization of fabrication tools such as 3d-printers, laser cutters, CNC machines, open source microcontrollers and open source software. Although many pieces of common laboratory equipment have been developed, software control of these items is sometimes lacking. Specifically, control software that can be easily implemented and enable user-input and control over multiple platforms (PC, smartphone, web, etc.). The aim of this proof-of-principle study was to develop and implement software for the control of a low-cost, 3d-printed microscope. Here, we present two approaches, which enable microscope control by exploiting the functionality of the social media platform Twitter or player actions inside of the videogame Minecraft. The microscope was constructed from a modified web-camera and implemented on a Raspberry Pi computer. Four aspects of microscope control were tested, including single image capture, focus control and time-lapse imaging. The Twitter-embodiment enabled users to send “tweets” directly to the microscope. Image data acquired by the microscope was then returned to the user through a Twitter reply and stored permanently on the photo-sharing platform Flickr, along with any relevant metadata. Local control of the microscope was also implemented by utilizing the video game Minecraft, in situations where Internet connectivity is not present or stable. A virtual laboratory was constructed inside the Minecraft world and player actions inside the laboratory were linked to specific microscope functions. Here, we present the methodology and results of these experiments and discuss possible limitations and future extensions of this work.

## INTRODUCTION

The general interest in using and developing low cost, open source labware is gaining considerable traction in garages, academic labs and commercial spaces [1]. This is largely being driven by the so-called “maker movement ” in which people are now exploiting the widespread popularity and accessibility of fabrication tools (3D printers, laser cutters, CNC machines, etc.) and open source electronics (Arduino, Raspberry Pi, etc) to build simple and advanced scientific equipment for a diversity of applications [2]. Designs and instructions are shared freely, typically under some form of open source license, and generally undergo several rounds of improvement as a result of the contributions from other users. Importantly, such innovations have matured to a level that it is possible to setup a basic, functional, laboratory space at extremely low cost. In addition, the spirit of “frugal science ” has led to several innovations in low cots diagnostic tools, often built around cell phone platforms. These approaches have the important potential for lowering the cost of diagnosis and treatment of diseases in both developed and developing countries.

In the context of the biological sciences, the “do-it-yourself biology (DIYBio) movement” has driven the development of several tools that are critical in any cell/molecular biology laboratory [3]. This includes open source pipettes [4], centrifuges [5], water baths [6], stirrers and hot plates [7], shakers [8], electrophoresis kits [9], incubators [10], PCR [11], [12] and qPCR [13] machines, and low cost kits for manipulating DNA or transforming bacteria [14]. One other key piece of equipment is a light microscope. Several designs and approaches have been developed for creating low cost light microscopes with reasonable magnification [15]. Such designs have resulted in microscopes that can operate in a variety of modalities including bright field, dark field and fluorescence [15]. Cellphone based microscopes have been developed in which the phone’s camera is simply employed as the imaging sensor. These approaches either mount a low cost set of optics directly to the cellphone camera [16] or mount the cell phone onto a simplified microscope stand that employs microscope objectives [17]. Alternatively, discarded webcams can be converted into the microscope by taking apart the camera and simply flipping the lens in front of the imaging sensor [18], or placing a low cost ball lens in front of an imaging sensor or the eye [19]. Lenses can also be sourced from a discarded CD-ROM drive or an optical mouse [20], [21]. The ability to generate functional, low cost microscopes is made more attractive as they can be simply produced from discarded electronics. Indeed, the simplest embodiment of a DIY Microscope can be achieved by simply placing a water drop on the cellphone front facing camera. These various approaches are not only important for educational purposes, but also have a significant role to play in developing low cost diagnostic tools for the lab or the field [4]–[21].

As low cost imaging tools (and general labware) become more prevalent there is an increasing need for the development of software control and monitoring solutions. In order to employ a DIY Microscope in a research setting, one may desire the ability to conduct imaging experiments without having to be physically present at the microscope in order initiate image capture. For example, conducting time-lapse experiments of cell growth on a microscope has been placed inside of a sterile, temperature and atmospherically controlled incubator. Therefore, the purpose and objective of this study was to develop a general proof-of-principle approach and physical embodiment of DIY Microscope control that relies on freely available programs that can be installed on a PC or cellphone. In order to achieve these goals, we first constructed a basic DIY Microscope by modifying the popular Rapsberry Pi (RPi) camera to act as an objective [18]. A 3D printed case for the RPi computer and camera was designed and contructuted to form a microscope stand. Finally, an DVD-ROM drive was used to create a moveable sample stage, allowing for focus control. Sample positioning along the optical axis of the microscope and image capture was controlled by a python script.

Once the DIY microscope was constructed, we developed user interfaces by exploiting three popular existing applications and their available application program interfaces (APIs). Here, we demonstrate the ability to use the popular Twitter interface to send commands to an Internet connected DIY microscope. We also implemented online data storage by uploading captured images, along with important metadata, to the photo-sharing network Flickr. In this scenario, the DIY Microscope was assigned a Twitter account (@DIYMicroscope), which monitored for simple commands sent by any other Twitter user. Simple message syntax was developed in order to allow other user to adjust microscope focus, capture single images or initiate time-lapse recordings. Upon image capture, all data was stored on a publically accessible Flickr account. In a second embodiment, we developed an approach to control a local DIY Microscope by exploiting the API of the popular video game Minecraft. Here, we first constructed a virtual lab inside of the Minecraft world. In this scenario one is able to “play ” within this virtual world and use gaming actions to control a physical DIY microscope. Simple actions inside of the videogame allowed one to again adjust microscope focus, capture single images or initiate time-lapse recordings. In this case, the images are stored locally on the RPi hard drive. In this embodiment, there is no need for a consistent or reliable Internet connection.

## MATERIALS AND METHODS

### DIY Microscope

A basic DIY Microscope was constructed by employing strategies that have been previously demonstrated [15], [18], [20]. Briefly, the microscope was constructed using a Raspberry Pi (RPi) model B+ as the control computer, an RPi camera module (Rasperry Pi foundation), a discarded computer DVD-ROM drive and a 3D printed frame (Makerbot Replicator 2) (Fig. 1A). All 3D printer files are available online at http://www.thingiverse.com/pellinglab. The original lens from the RPi camera module was removed prior to installation, leaving only the image sensor (OmniVision OV5647-5MPx). A web camera lens (Logitech c310) was then inverted and installed on top of the image sensor. The RPi camera module and the lens were maintained in fixed positions relative to one another by mounting in a 3D printed mount.

The DVD-Rom drive was then disassembled leaving only the stepper motor, the laser pickup assembly and the frame intact. The drive was then mounted perpendicular to the RPi camera assembly in order to create a sample positioning stage that could be moved along the optical axis of the microscope in an inverted configuration. In order to fix the positions of each component of the entire assembly, a 3D printed frame was produced to which all components could be mounted. Finally, a sample stage was 3D printed and mounted to the laser pickup assembly. This configuration allowed us to easily adjust sample position and focus under computer control. The movement of the sample tray was achieved by controlling the movement of the stepper motor by employing the Easy Driver Stepper Motor Driver V4.4 (http://www.schmalzhaus.com/EasyDriver) in 8-step microstepping mode. The driver and a standard 5mm white LED were connected to RPi via GPIO pins in order to control sample positioning and illumination (Fig. 1B).

To calibrate the DIY microscope, we acquired images of standard microfabricated cantilevers commonly employed used in atomic force microscopy. The cantilevers have known dimensions. The larger cantilever in the image is 200 μm, corresponding to 190 pixels in a 1600 by 900 pixel image (Fig. 1C).

**Figure 1.**
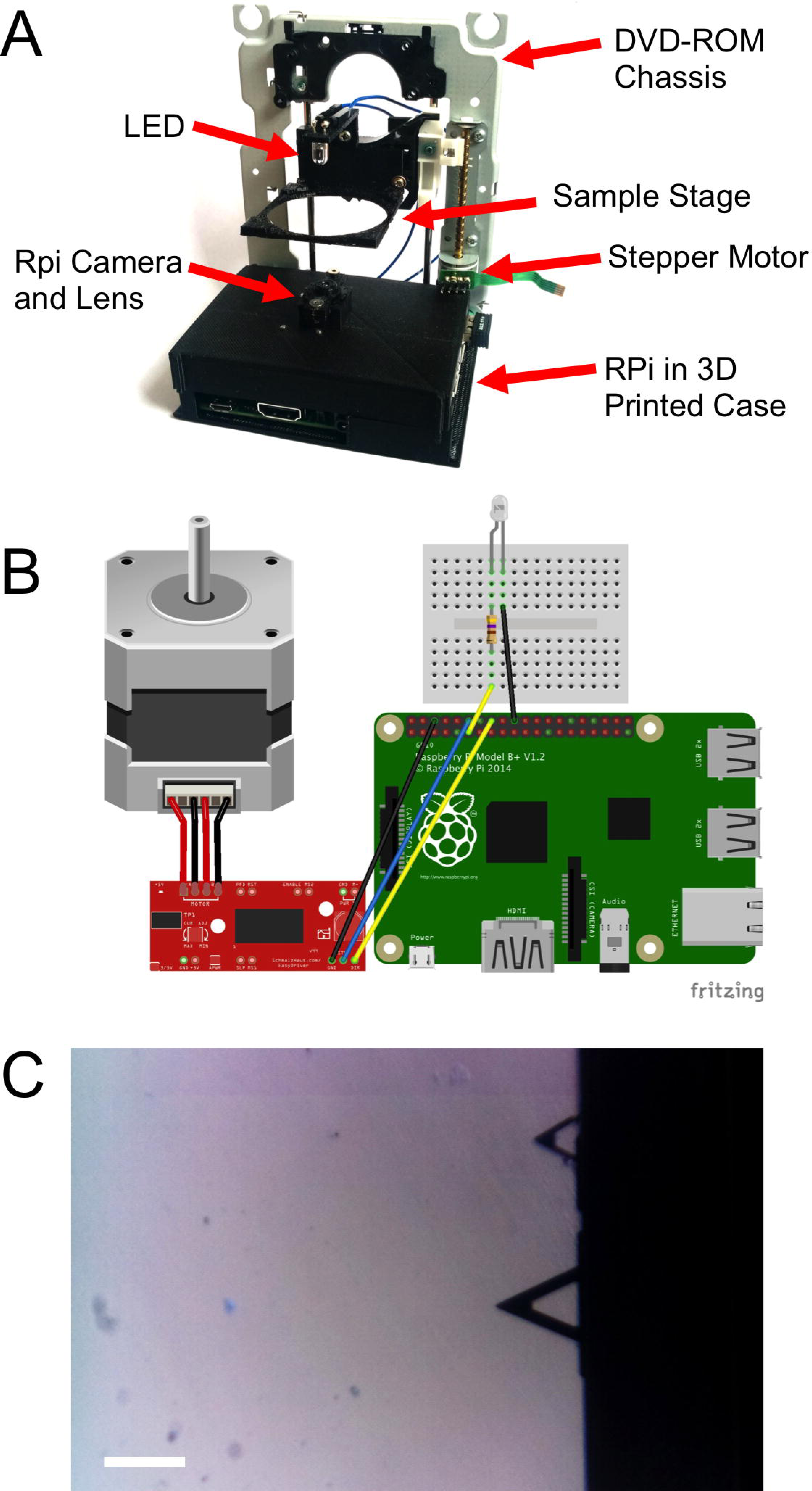
DIY microscope construction. A) A case was 3D printed in order to mount the RPi computer, camera and lens assembly and DVD-ROM drive chassis. A sample stage was also 3D printed and mounted to the laser pickup assembly. B) A simple motor driver was then employed to control the stepper motor with the GPIO pins of the RPi. C) Calibration of the microscope was achieved by acquiring images of microfabricated atomic force microscopy cantilevers. The lower cantilever has a known length of 200 μm, corresponding to 190 pixels in the image (scale bar = 200 μm).

### Online Control of the DIY Microscope

A python script was written that allows any Twitter subscriber to remotely interact with the microscope via the Twitter app or website. The code we developed is available online at https://github.com/pellinglab. The open source python library Tweepy (tweepy.org) was employed to facilitate communication with Twitter’s application programming interface (API). The microscope was assigned the Twitter account @DIYMicroscope in order to facilitate user interaction. The python script running on the RPi monitors‘tweets’sent to the account @DIYMicroscope and examines them for simple key words. For example, the Twitter user can capture single images, control sample positioning and focus, and initiate time-lapse imaging (details are presented in the Results and Discussion section).

### Online Image Capture and Storage

The RPi storage capacity can be limited as it is defined by the capacity of the SD card the user has employed. In order to prevent memory issues associated with a large number of image acquisitions, we utilized the photo-sharing website Flasickr (flickr.com), as an image hosting platform. To remotely and automatically interact with the Flickr API, we implemented the Beej Flickr API python library (http://stuvel.eu/flickrapi). Each time an image is acquired by the script (i.e a single frame or an concatenate of frames), a copy is uploaded to the DIY microscope’s Flickr account. Then the original image is removed then from the RPi hard drive to save space.

### Offline Control of the DIY Microscope

To locally control the microscope, we designed a python script that will create a user interface in the videogame universe of Minecraft (Mojang). The code we developed is available online at https://github.com/pellinglab. We constructed a virtual lab in which user actions during game play can be used to initiate specific microscope functions. To achieve this, we employed the Minecraft python API library (http://www.stuffaboutcode.com/p/minecraft-api-reference.html). Inside the virtual laboratory, the a sword-equipped player can control the microscope by performing specific actions with ‘control blocks’ inside the virtual laboratory. Block actions were designed to allow the user to generate a live preview, capture an image and adjust the stage position for focus control. Captured images are stored in a specific local folder for future use.

## RESULTS AND DISCUSSION

### Software Design for Twitter Control

When the python script is launched, an authentication procedure to *Flickr* is initiated (a *Yahoo*! account, API key and API secret are required), followed by an authentication to twitter (a *Twitter*account, API key and API secret, token and secret token are required). Both accounts must be established and verified by the system administrator in advance. The Flickr account is required for storage of images acquired by the DIY Microscope. The DIY Microscope will be addressed and controlled by sending ‘tweets’ that mention the system-specified Twitter account. When the authentication between Flickr and Twitter is correctly established, the script connects to Twitter’s streaming API with specific keywords. This allows the program to obtain real-time tweets from the social network. When a user sends a tweet to the system-specified Twitter account containing single, or multiple, keywords, a JavaScript Object Notation (JSON) object is returned by the streaming API, containing parameters such as the user name, screen name, location, *tweet* content and the time. The JSON object is then examined and conditional actions are undertaken depending on the keywords identified in the tweet (Fig. 2). Importantly, to avoid any unwanted interactions with the DIY Microscope (i.e. a random user has one of the keywords in their tweets), the requesting user must include the system-specified twitter handle (e.g. @example) in their tweet.

**Figure 2.**
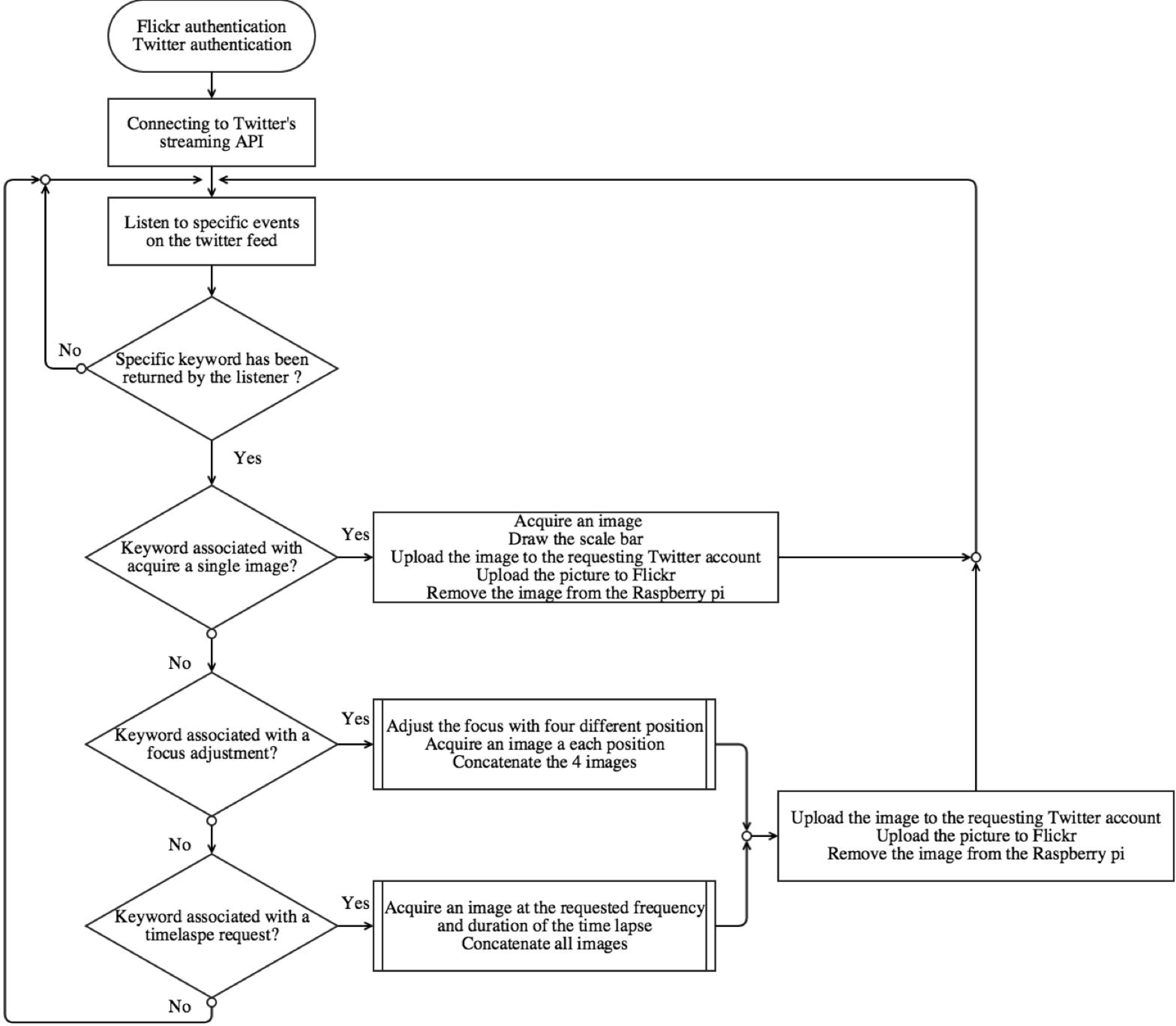
Flow-chart representing our python script that enables Twitter-based user interaction with the DIY Microscope

In this embodiment, we designed four types of user interaction, such as taking a single image, initiating a timelapse, adjusting the focus and obtaining a ‘focus group’ image (Fig. 3 and Fig. 4). When the tweet contains the keyword ‘singleimage’(Fig. 3A), the current image frame captured by the RPi camera is temporarily stored on the RPi, a scale bar is drawn on the image and returned to the requesting user in a Twitter message (Fig. 3B). In addition to the returned image, the reply tweet also includes the message ‘@user scale bar is X_UNITS”, where @user is the Twitter handle of the requesting user and X_UNITS = value determined by the user after microscope calibration. Finally, the temporarily stored local copy of the image is then uploaded to the user specified Flickr account and permanently removed from the RPi. In Fig. 3, a single image is requested by the user @pellinglab and an image of cellulose derived from apples [22] is acquired and returned by the microscope.

**Figure 3.**
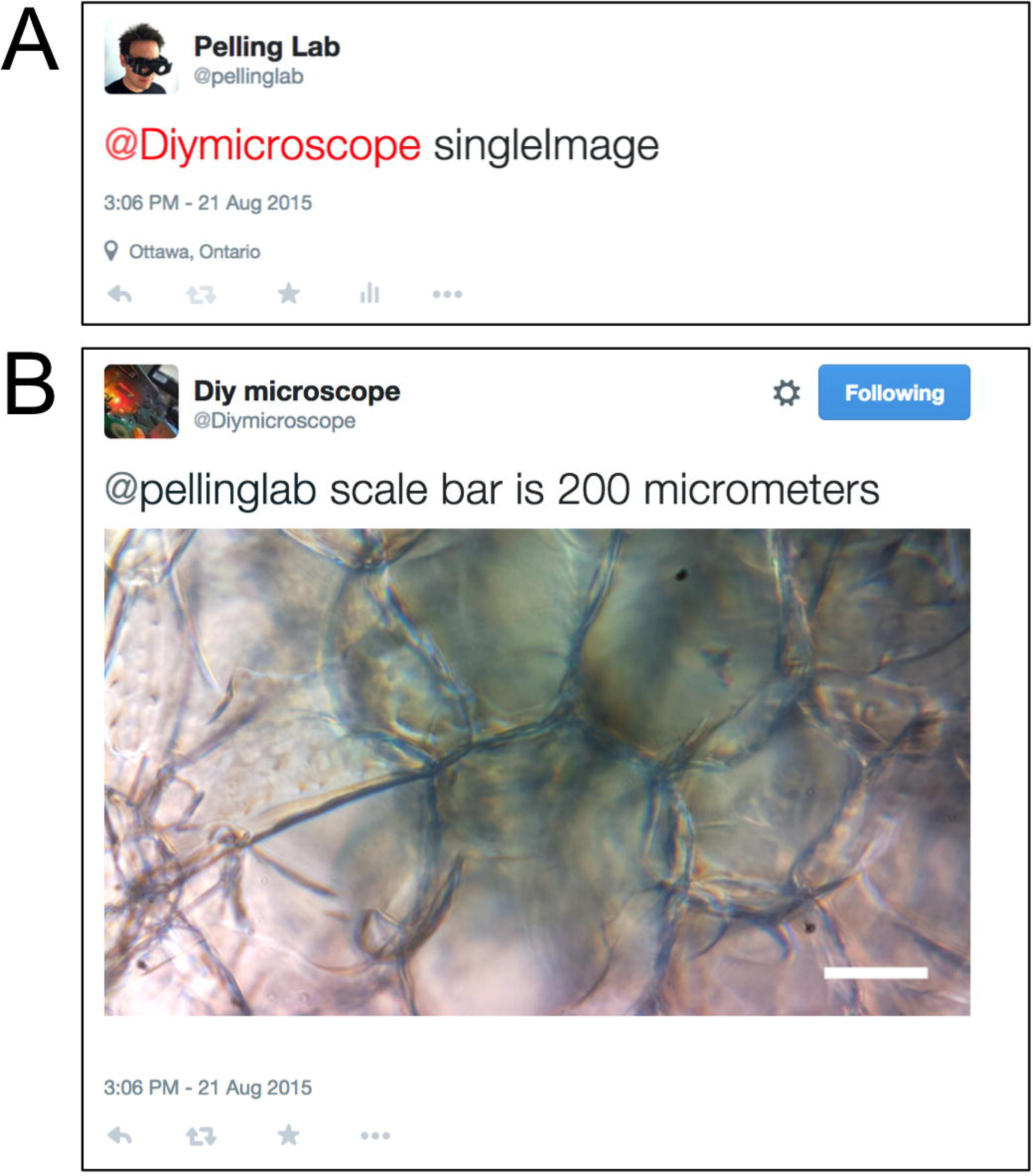
Twitter acquisition of a single image from the DIY Microscope. A) The account @pellinglab initiates image acquisition by posting a tweet that mentions the account @DIYMicroscope and includes the keyword‘singleImage’. Other text can be included in the message but the script will ignore them. B) The DIY microscope acquires an image and responds to the resting account.

**Figure 4.**
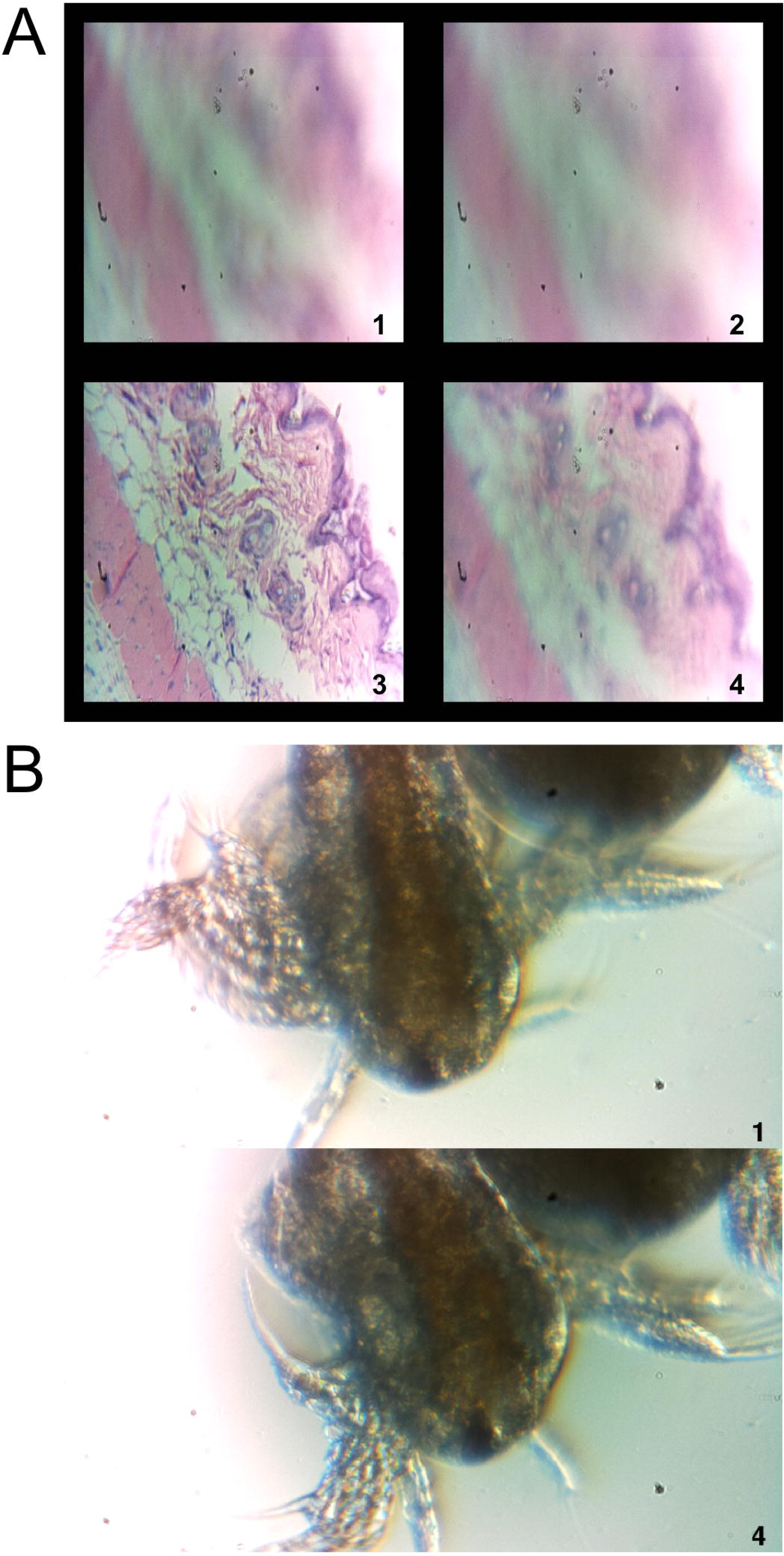
Focus and time-lapse control through Twitter interactions. A) The command‘diyfocus dofocus N’initiates image capture of the sample when moved to four specific positions. The user is then provided with a concatenated image containing the four acquired, and indexed, images. This routine allows the user to determine the optimal sample position in order to set the focus. In this case, a sample histology slide was employed for imaging. As these images are only for stage positioning purposes no scale bar is included, however each image is 0.95 × 0.95 mm. B) It is also possible to initiate time-lapse imaging using the keywords ‘diytimelaspe duration D frequency F’, D and F, correspond to integer values with units of minutes and frames/minute, respectively. In this case, brine shrimp (*Artemia*– commonly grown as live feed for fish larva) were imaged with the microscope. A final image is returned to the user that only contains the first and last image acquired during the time−lapse interval. All images from the sequence are stored on Flickr along with any relevant metadata. No scale bar is provided in this example, however the scale is known after user calibration and relevant details are included in the Flickr metadata. In this case each image is 1.70x0.95millimeters.

To move the stage in order to focus the sample, the user sends a tweet to the DIY Microscope that includes the keywords ‘diyfocus further N’ or ‘diyfocus closer N’(N = integer value). The sample stage will then move further away, or closer to, the RPi camera by N microsteps. An image will be acquired after moving to the new position, sent back to the user through twitter and stored on Flickr as above. If the user wants to sample multiple focus positions, a tweet is constructed which includes the keywords‘diyfocus dofocus N’ (N = integer value). In this scenario, the sample platform is moved away from the RPi camera by N microsteps and an image is acquired. The sample then moves another N microsteps away from the camera a second image is obtained before moving back to the original position. The process is then repeated in the opposite direction in order to acquire image number 3 and 4. The four locally stored images are uploaded to Flickr, along with their metadata (the twitter handle of the requesting user, the original message sent by the user, N and the corresponding frame number), and then concatenated into a single 900 by 900 pixel image (Fig. 4A). The concatenated image is then returned to the user with the corresponding frame numbers printed on each sub-image along with the message ‘@user your N microstep dofocus sequence is complete’ (where @user and N are determined from the original user message). The user can now adjust the stage position to the desired location (using the‘diyfocus further/closer’ keywords, followed by an appropriate integer value). In Fig. 4A, the concatenated image of a hematoxylin and eosin stained histology sample being moved in and out of focus is shown.

Finally, the user can initiate a time lapse using the ‘diytimelapse duration D frequency F’ keywords, replacing the ‘D’ and ‘F’ with integers for duration and frequency, respectively. D and F, values assume units of minutes and frames/minute, respectively. The timelapse is carried out as requested and each acquired image is uploaded to Flickr along with its corresponding metadata (the twitter handle of the requesting user, the original message sent by the user, D, F and the corresponding frame number). Upon completion of the timelapse, a concatenated image containing the first and last frame of the time lapse is constructed and returned to the user with the frame numbers printed on each sub-image (Fig. 4B). The returned image also includes the message ‘@user your timelapse of D minutes at F frames/minute is now complete’ (where @user, D and F are determined from the original user message). In Fig. 4B, time-lapse microscopy was conducted on a sample of *Artemia* (brine shrimp) and the image sent back to the user through twitter is shown. Four images were acquired (duration of one minute at a frequency of four images per minute) and as described above, only the first and last images are sent to the user through twitter. All images in the sequence are stored on Flickr for later analysis by the user.

Importantly, whenever an image is uploaded to Flickr (irrespective of the keyword(s) employed), specific metadata is included with the image in order to allow for identification and filtering. The metadata associated with each image includes the twitter handle of the requesting user, the keyword(s), a timestamp and the frame number.

### Software Design for Minecraft Control

In situations where a user may lack Internet access, we designed an approach for local, offline, control of the DIY Microscope. In this case, we designed a python script that allows for user interaction through the videogame Minecraft. Conveniently, Minecraft is already included in the freely available Raspbian distribution of Linux for the RPi. When our python script is launched, the player position is immediately updated to be facing the virtual ‘laboratory’ (Fig. 5).

**Figure 5.**
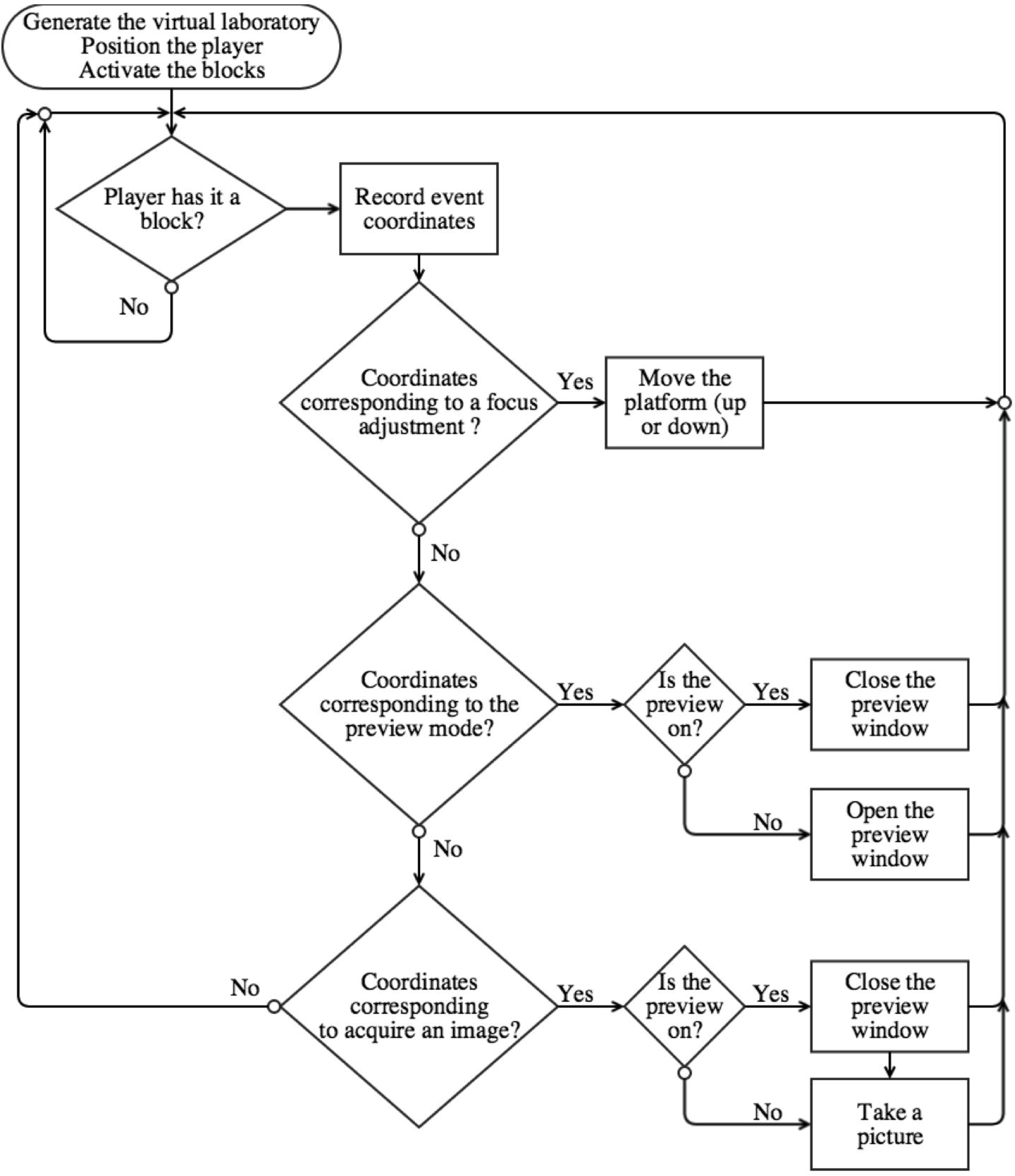
Flow−chart representing our python script that enables Minecraft−based user interaction with the DIY Microscope

Upon entering the laboratory, a sword-equipped player can now interact with the DIY microscope ‘control blocks’ (Fig. 6A). When standing in front of the desired control block, the right mouse button can be used to initiate a specific action. Such actions will return an event object, containing the coordinates of the specific block as a tuple. The coordinate tuple is then used to initiate conditional actions that are used to control the ‘real-life’ DIY Microscope. When the player activates one the extremity blocks (black and grey blocks), the stepper motor will perform N microsteps clockwise (or counterclockwise). The user can specify N in the python script. The yellow block will display a live preview of the sample, on a 640 by 480 pixels window (Fig. 6B). The blue blocks will take a single image when activated, and save the image as a “.png ” file on the RPi. To avoid over usage of the GPU memory, the program will close live preview (if open) prior to acquiring an image. A message on the minecraft user-interface is displayed when the image is successfully stored (Fig. 6C).

**Figure 6.**
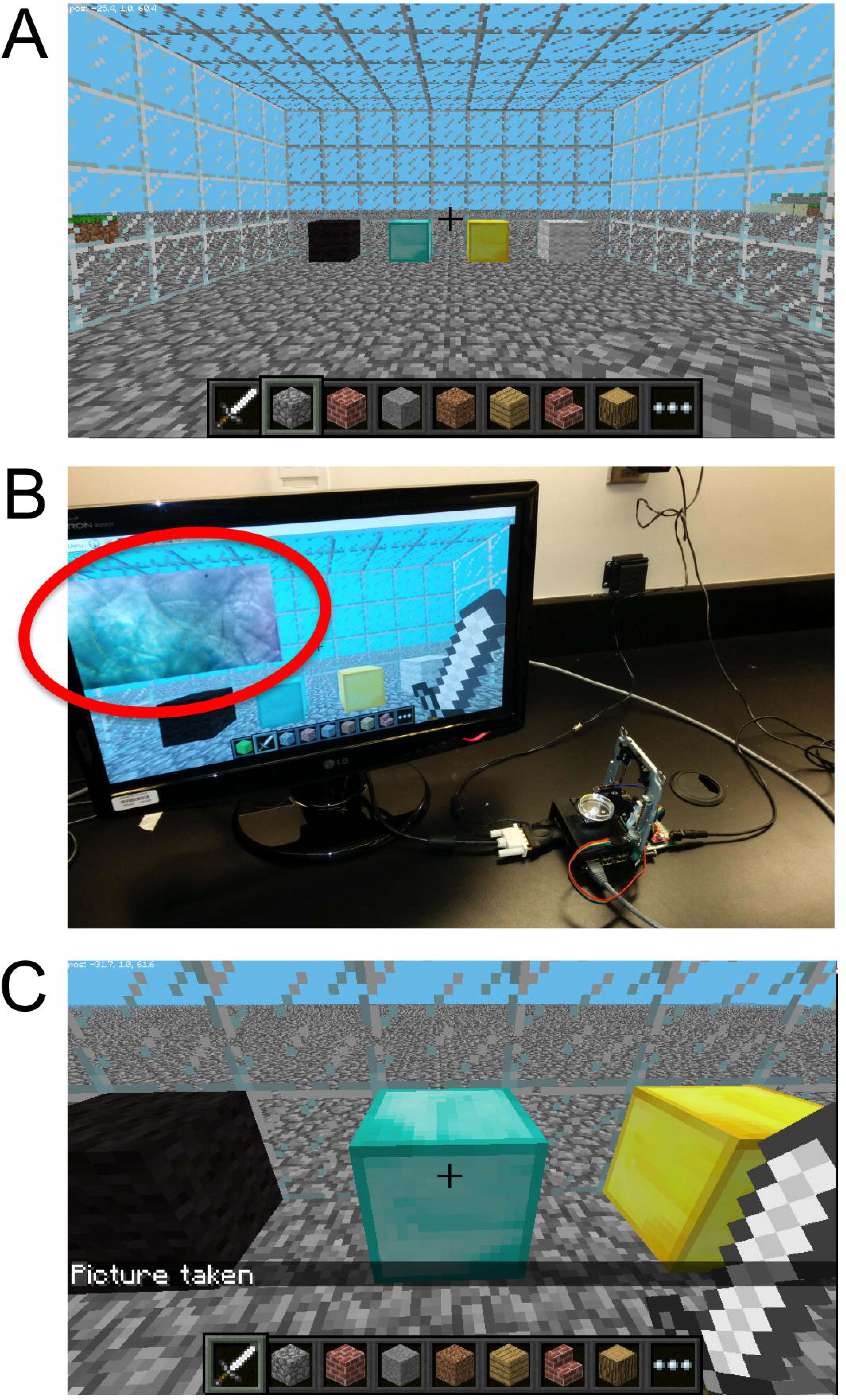
The Minecraft environment. A) The user simply uses the sword to control a locally connected DIY Microscope by interacting with the four control blocks inside of a virtual ‘laboratory’. The blocks allow the user to adjust the sample stage up and down (black/gray blocks), B) obtain a live preview (yellow block) and C) acquire an image (cyan block).

## CONCLUSIONS

### Possible Limitations

The Twitter public streaming API limits the application to a fixed number of keyword filters per application [23]. The current application only requires 3 keywords (“singleImage”,“diyfocus” and “diytimelapse”) to initiate user-winteraction, however, more complex interactions may require many more keywords. As well, user-interaction is also limited by the number of allowed connections to the API per hour [23] but the exact number is not currently publically available. Importantly, a user will also receive an error message from Twitter if they post the exact same tweet multiple times over a short time period. Therefore, Twitter will not allow a single user to post the tweet ‘@DIYMicrosocpe singleImage’ more than once. The user can overcome this limitation by adding more text to the tweet or deleting the original tweet. Currently, the Flickr API also limits the application to 3600 requests per hour [24].

Some other limitations can also arise if multiple users are attempting to interact with the microscope simultaneously. For instance, if a time-lapse sequence is initiated, the script will complete the image acquisition before returning to the stream listener. In this case, a user sending a request will not get an immediate response from the microscope.

Finally, the microscope is inherently limited to the presence and stability of Internet connectivity available to the RPi. In the case of a lost connection, the script will have to be re-executed in order to establish a connection to the Twitter streaming API and Flickr API. To overcome the issue with Internet connectivity, we also implemented a control interface within the Minecraft universe. Of course, electrical power is always required as with any modern microscope. However, it is possible to power the RPi using solar panels and batteries in cases where microscopic imagery is required but electrical connections are not easily accessed. As the Minecraft implementation of the DIY Microscope stores pictures locally, the maximum storage space will depend on the SD card capacity. Cloud storage can be implemented with the *Minecraft* program, but will require an Internet connection. However, if storage space becomes a limitation, the user can move the files to an alternative storage device (usb stick or via local file transfer) or employ a larger SD card.

### Possible Extensions

Future versions of both the *Twitter* and *Minecraft* interfaces could include image analysis features, such as cell counting, cell tracking for time lapses and thresholding by implementing OpenCV [25]. Such integration could allow the user to obtain qualitative and quantitative from the sample, in a remote or local way. An automatic focusing library could also be implemented to both program to enhance image quality acquisition rapidity. In its current implementation, our approach does not maintain privacy as all micro blogging posts and pictures on *Twitter* and *Flickr* are publicly available. In order to overcome this potential limitation, a smart phone based application could easily be developed to interact with the current version of the microscope, either via a direct connection (*wifidirect* or *bluethooth*) or a web−based server. Such a configuration could allow the user to use customize features, independent from *Twitter, Flickr* and *Minecraft APIs*, consequently avoiding the rate limitation from the third-parties. Of potential interesti is that both of our approaches can also be exploited on other types of laboratory equipment. One can easily interface the multiple outputs of the RPi to an existing or “hand−made ” equipment, or extends existing programs [1]–[3]. Future use of the *Minecraft* program can let access to multiple player in the “virual laboratory”, either by a local or internet connection. This let the possibility for the “players” to cooperatively interact and controls “real” physical laboratory equipment inside a virtual world.

## ACKNOWLEDGMENTS

The authors would like to thank Daniel Modulevsky and Dr. Charles Cuerrier for providing the histological sample for imaging.

